# Distinct Cellular Strategies Determine Sensitivity to Mild Drought of Arabidopsis Natural Accessions

**DOI:** 10.1101/2020.08.24.265041

**Authors:** Ying Chen, Marieke Dubois, Mattias Vermeersch, Dirk Inzé, Hannes Vanhaeren

## Abstract

The world-wide distribution of *Arabidopsis thaliana* (Arabidopsis) accessions imposes different types of evolutionary pressures, which contributes to various responses of these accessions to environmental stresses. Drought stress responses have been well studied, particularly in Columbia, a common Arabidopsis accession. However, the reactions to drought stress are complex and our understanding of which of these responses contribute to the plant’s tolerance to mild drought is very limited. Here, we studied the mechanisms by which natural accessions react to mild drought at a physiological and molecular level during early leaf development. We documented variations in mild drought tolerance among natural accessions and used transcriptome sequencing of a drought-sensitive accession, ICE163, and a drought-tolerant accession, Yeg-1, to get insights into the mechanisms underlying this tolerance. This revealed that ICE163 preferentially induces jasmonates and anthocyanin-related pathways, which are beneficial in biotic stress defense, while Yeg-1 has a more pronounced activation of abscisic acid signaling, the classical abiotic stress response. Related physiological traits, including content of proline, anthocyanins and ROS, stomatal closure and cellular leaf parameters, were investigated and linked to the transcriptional responses. We conclude that most of these processes constitute general drought response mechanisms that are regulated similarly in drought-tolerant and -sensitive accessions. However, the capacity to close stomata and maintain cell expansion under mild drought appeared to be major factors that contribute to a better leaf growth under mild drought.

**One-sentence summary:** This paper demonstrates that an efficient closure of stomata and maintenance of cell expansion during drought conditions are crucial to maximally preserve plant growth during water deficit.

## INTRODUCTION

With the increasing effects of global warming and climate change, agricultural crop production is facing major challenges. Drought is one of the main abiotic stresses throughout the world and limits agricultural yield by reducing plant growth (Araus et al., 2002). Drought stress can occur in a broad spectrum, ranging from mild drought, which leads to growth arrest, to severe drought in which plants suffer from desiccation and wilting, eventually leading to their death. In the previous decades, numerous studies have focused on severe drought stress. However, mild drought stress is occurring more often in actual field conditions. Recent studies have demonstrated that plants apply different regulatory strategies to cope with mild drought compared to severe drought stress (Harb et al., 2010; Skirycz et al., 2011; Ma et al., 2014; Clauw et al., 2015). For example, only one third of differentially expressed genes overlapped between severe drought and mild drought treatments in young developing Arabidopsis leaves (Clauw et al., 2015) and plants that were described to be tolerant to severe drought stress did not perform better under mild drought conditions (Skirycz et al., 2011). Exploring the molecular and physiological strategies that plants apply to cope with mild drought stress is therefore essential to ensure our future agricultural productivity.

Being sessile, plants have evolved complex regulatory strategies for transcription, metabolism, and morphology to overcome drought stress (Claeys and Inzé, 2013; Tardieu et al., 2014). Under water-deficient conditions, plants limit their shoot growth to reduce water losses. The development of leaves, which is strictly controlled by cell proliferation and cell expansion (Gonzalez et al., 2012), can be significantly and rapidly affected by drought stress (Skirycz and Inzé, 2010). Especially, water deficit greatly suppresses cell expansion due to the low turgor pressure (Feng et al., 2016). Because leaves are the plant source tissues through photosynthesis, reductions in their size will lead to a significant yield penalty.

Phytohormones play an important role in responses to drought stress. The most well-known drought-related hormone, abscisic acid (ABA), accumulates rapidly during drought conditions (Christmann et al., 2005). The mechanisms of ABA-mediated drought stress responses have been well studied (for a review, see Nakashima and Yamaguchi-Shinozaki, 2013; Yoshida et al., 2014). During drought stress, several key elements of the ABA signaling pathway, such as *SNF1-RELATED PROTEIN KINASES 2* (*SnRK2s*) (Boudsocq et al., 2004), *ABSCISIC ACID-RESPONSIVE ELEMENT BINDING PROTEIN 1* (*AREB*1) and *AREB2* (Fujita et al., 2005) are upregulated. In addition, ABA accumulation by water deficits leads to the rapid closure of stomata (Israelsson et al., 2006), which increases water use efficiency and, hence, results in an improved survival during drought stress. On the other hand, jasmonates (JAs) and ethylene (ET), which are known to play essential roles in orchestrating plant defense against pests and pathogens, also influence abiotic stress tolerance (for a review, see Kazan, 2015). Under water-deficient conditions, JA interacts with redox processes and enhances the synthesis of antioxidative stress compounds, such as ascorbate and glutathione (Shan and Liang, 2010). Moreover, transcriptome studies have shown that JA-related genes are the most relevant class of non-ABA genes that are responsive to severe drought stress (Huang et al., 2008; Harb et al., 2010). The effect of ET in drought stress is bidirectional. During drought stress, ET inhibits ABA-induced stomatal closure (Tanaka et al., 2005). On the other hand, ET promotes stomatal closure by prompting NADPH oxidase-mediated production of scavenging reactive oxygen species (ROS) in stomatal guard cells (Desikan et al., 2006).

Some metabolites, such as anthocyanins and proline, also act in the defense against drought (Bhaskara et al., 2015; Li et al., 2017). Anthocyanins are synthesized via the phenylpropanoid pathway in plants and have been described to play an important role in biotic and abiotic stress tolerance by absorbing ultra violet (UV), high light irradiation and ROS (Landry et al., 1995; Mittler et al., 2004). During drought stress, increased biosynthesis and decreased catabolism result in a higher accumulation of proline, limiting the retardation of growth (Kishor et al., 1995). Proline counteracts osmotic stress in several aspects: as an osmolyte (Handa et al., 1986), as a stabilizer of protein structures (Wu and Bolen, 2006), as a scavenger of free radicals (Smirnoff and Cumbes, 1989), as a sink for energy (Hare and Cress, 1997) and by maintaining the NADP^+^/NADPH balance (Sharma et al., 2011). Recent research revealed a dynamic and accession-dependent proline accumulation under drought stress (Kesari et al., 2012; Bhaskara et al., 2015).

Adverse conditions can also trigger the production of ROS in plants, including superoxide anion radicals (O_2_ ^-^), hydroxyl radicals (OH•), hydrogen peroxide (H_2_O_2_) and singlet oxygen (O^1^_2_). ROS can act as signal transduction molecules in plants, but a high accumulation of ROS will lead to severe cellular damage, the inhibition of photosynthesis and even plant death (Dat et al., 2000). ROS accumulation during stress largely depends on the balance between ROS production and ROS scavenging (Mittler et al., 2004). Superoxide dismutases (SODs) act as primary enzymes in ROS scavenging. Overexpression of SOD-encoding genes can enhance the tolerance of plants to severe drought (Liu et al., 2013). In addition, the ROS balance in plants under stress conditions can also be regulated by JAs (Miller et al., 2010), anthocyanins (Li et al., 2017) and proline (Liu et al., 2015).

These different, interconnected stress response strategies provide a broad regulatory potential to preserve plant fitness during unfavorable conditions (Pieterse et al., 2009; Berens et al., 2019). While all of these responses have most often been studied in the context of moderate to severe drought that was applied to adult plants, little is known about how plants that originate from different natural environments employ and coordinate their growth to mild drought stress. Arabidopsis has been used as a model for plant research for a long time and many natural accessions are widely distributed in various geographical areas. These diverse natural habitats each exert different evolutionary pressures, which allowed the accessions to develop different molecular and physiological strategies to adapt to their specific environment (Weigel, 2012). Hence, specific strategies to cope with water deficits between natural accessions can be expected. Exploring the specific responses of natural accessions to mild drought can therefore give more insight into the diverse regulatory networks, which can be useful for plant bleeders. Here, we compared the response of fifteen natural accessions to mild drought and found that the leaf growth of these ecotypes responds in a broad range of sensitivity. To identify the discriminating factors that underlie this phenotypic difference response, we explored the transcriptome and physiological changes of tolerant and sensitive accessions after a mild drought treatment and identified different transcriptional responses. We found that the efficiency in stomatal closure and especially the capacity to maintain cell expansion during drought are crucial to preserve organ growth.

## RESULTS

### Natural Arabidopsis Accessions Reduce Growth to a Different Extent under Mild Drought

To explore how genetic diversity in Arabidopsis can affect the response to mild drought stress, we screened the growth of fifteen natural accessions from different origins (Fig. 1A, Supplemental Table S1) on the automated Weighing, Imaging and Watering Machine (WIWAM) (Skirycz et al., 2011). The mild drought (MD) treatment was initiated for half of the plants at 6 days after stratification (DAS), when the third true leaf (L3) starts to emerge. The other half of the plants was kept under well-watered (WW) conditions to serve as a control (for the experimental details, see Material and Methods). The plants were harvested at 22 DAS and the area of the mature L3 was measured. The average leaf area (LA) already differed between the accessions under WW conditions (Fig. 1), but except for EY15-2, all accessions showed a significant relative reduction in LA under MD compared to WW conditions (Fig. 1B). Remarkably, the reduction in LA was very different depending on the accession, ranging from 14% to 61% (Fig. 1B, Supplemental Table S2). The sensitivity to MD (% reduction) did not depend on the size of the leaf under WW conditions, since no correlation between LA under WW conditions and relative reduction by MD could be observed (Supplemental Fig. S1). We identified drought-sensitive accessions, such as Oy-0, Ler-0, ICE97 and ICE163, as well as more tolerant accessions, including C24, Yeg-1, An-1, Sha and EY15-2.

**Figure 1.**
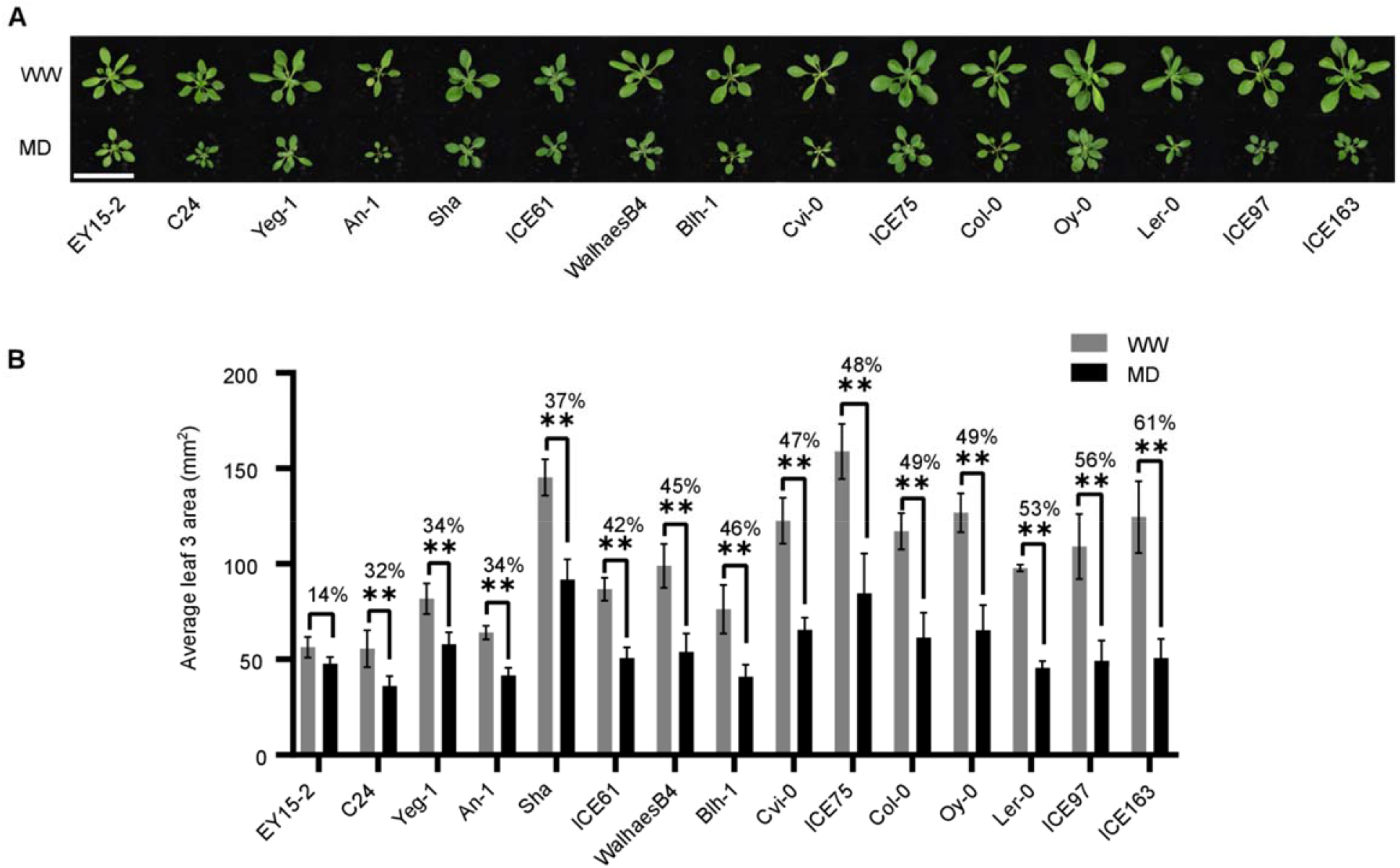
Different Arabidopsis accessions show a different leaf growth reduction under mild drought. **(A)** Rosettes of 15 natural accessions under well-watered (WW) and mild drought (MD) conditions at 22 days after stratification (DAS). Scale bar = 5 cm. **(B)** Average area of the third leaf under WW and MD conditions at 22 DAS. Error bars represent the SE, n = 3 biological repeats. Percentages represent relative reductions in leaf area under MD conditions compared to the WW control. Asterisks indicate a significant difference as determined by ANOVA, TukeyHSD. (** p adj <0.01). Significance of relative reduction in leaf area differences between accessions is shown in Supplemental Table S2 (* p adj < 0.05; ANOVA, Ecotype X Treatment interaction).

### Mild Drought Triggers Different Transcriptomic Changes in ICE163 and Yeg-1

To gain further insights into the molecular mechanisms that underlie the different leaf growth responses under MD stress, we selected two accessions that were significantly affected by drought, but at opposite ends of the spectrum. ICE163 and Yeg-1 were chosen as representatives of sensitive and tolerant accessions, respectively (Fig. 1B, Supplemental Table S2). Using the same drought setup as described above, we isolated the actively growing L3 of MD plants and the respective WW controls at five days after water retention (11 DAS) for transcriptome profiling by RNA-sequencing. A multidimensional scaling (MDS) plot showed that the samples clustered mainly by ecotype (Supplemental Fig. S2A), illustrating the profound effect of genetic background on the transcriptome. Even in absence of the stress treatment (WW conditions), ICE163 and Yeg-1 had very different basal gene expression levels. In total, 22.1% of all expressed genes were differentially expressed (DE) between both accessions, including 1853 and 2736 genes with higher expression in Yeg-1 and ICE163, respectively (Supplemental Fig. S2B).

Next, we compared the transcriptomic responses that were altered by MD treatment in ICE163 and Yeg-1. We identified 514 genes that were DE in ICE163 and 430 genes in Yeg-1 (MD vs. WW per accession, FDR ≤ 0.05, |log_2_FC|≥ 0.6). In total, 178 genes (143 upregulated, 35 downregulated) were in common between these accessions (Fig. 2, A-C). Within the common MD-induced genes, GO categories such as superoxide radicals (FDR = 0.000213), JA biosynthesis (FDR = 0.0252), cell wall modification (FDR = 0.035) and the ABA signaling pathway (FDR = 0.0171) were overrepresented (Supplemental Table S3). Overrepresented GO categories of the common downregulated genes included the response to ozone (FDR = 0.0102) and superoxide (FDR = 0.00313) (Supplemental Table S3).

**Figure 2.**
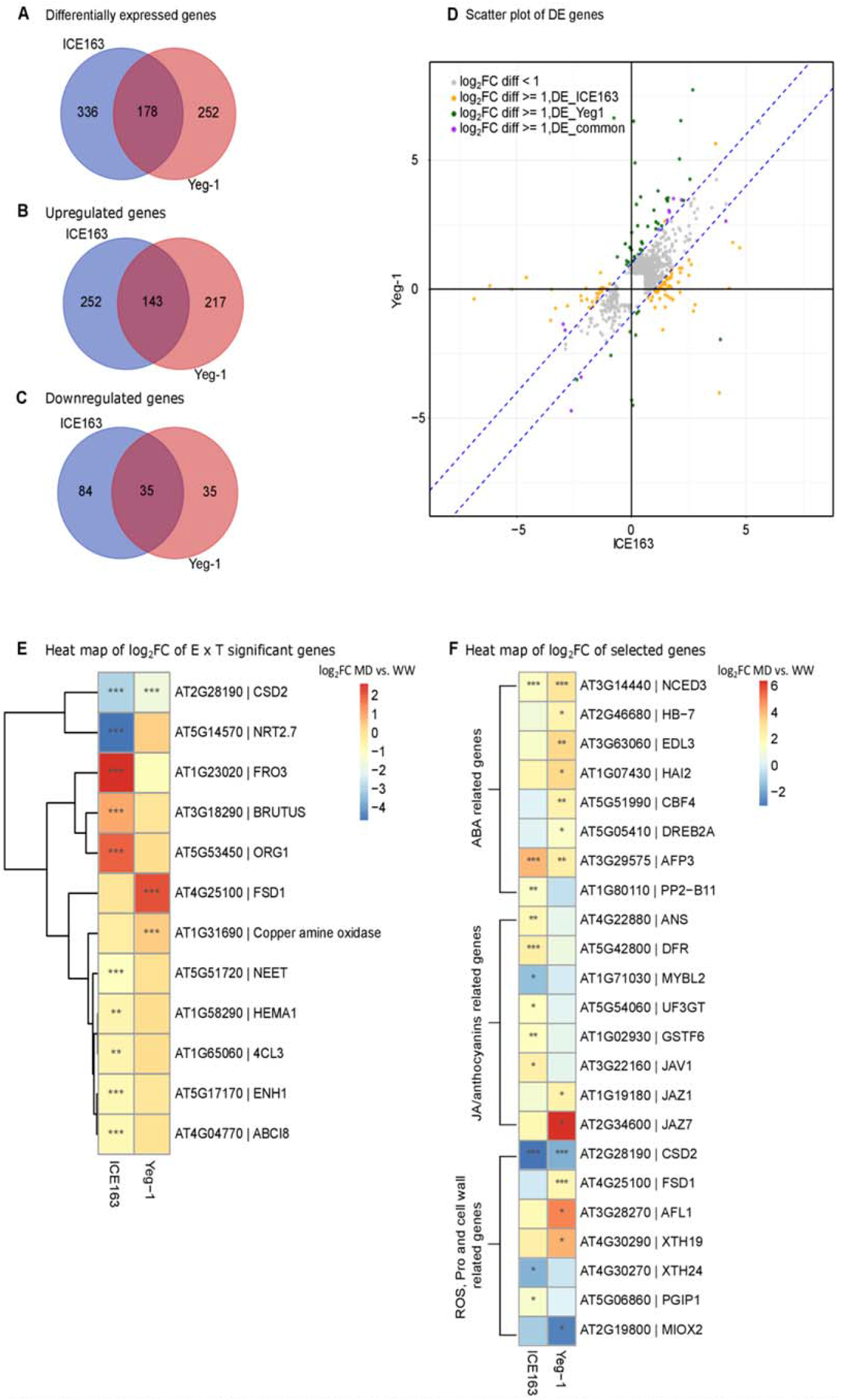
ICE163 and Yeg-1 react differently to mild drought at the transcriptome level. **(A)** Venn diagram of differentially expressed (DE) genes (MD vs. WW) in ICE163 and Yeg-1 (FDR < 0.05, log_2_FC ≥ 0.6 or log_2_FC ≤ -0.6). **(B)** Venn diagram of upregulated genes by MD (FDR ≤ 0.05, log_2_FC ≥ 0.6). **(C)** Venn diagram of downregulated genes by MD (FDR < 0.05, log_2_FC ≤ -0.6). **(D)** Scatter plot of DE genes in ICE163 and Yeg-1. Log_2_FC (MD vs. WW) of gene expression levels in Yeg-1 are plotted against those in ICE163. Gray: genes whose log_2_FC (MD vs. WW) difference between Yeg-1 and ICE163 is less than 1; orange: genes whose log_2_FC (MD vs. WW) difference between Yeg-1 and ICE163 is equal or larger than 1 and their expression is only significant in ICE163; green: genes whose log_2_FC (MD vs. WW) difference between Yeg-1 and ICE163 is equal or larger than 1 and their expression is only significant in Yeg-1; purple: genes whose log_2_FC (MD vs. WW) difference between Yeg-1 and ICE163 is equal or larger than 1 and their expression is significant in both Yeg-1 and ICE163. **(E)** Heat map of log_2_FC of genes that show a significant (FDR < 0.05) Ecotype x Treatment (E x T) interaction. Red and blue correspond to increased or decreased expression upon drought, respectively. **(F)** Heat map of the log_2_FC of selected genes from the E x T interaction and scatterplot. Red and blue correspond to increased and decreased expression (MD vs. WW), respectively. In D and F, asterisks indicate that the expression levels are significantly different from the control condition (WW) as determined by ANOVA, TukeyHSD. (* FDR < 0.05; ** FDR <0.01; *** FDR <0.001). For gene names and abbreviations, see Supplemental Table S8.

To search for transcriptional mechanisms that could underlie the different phenotypic responses of Yeg-1 and ICE163 to drought, we further explored the specific DE genes in these accessions. When looking at the genes that were DE in each accession during MD (Fig. 2A), we found that 336 and 252 genes were DE specifically in ICE163 or Yeg-1, respectively (Fig. 2A, Supplemental Table S4). However, we realized that some of these genes could actually be affected by MD to a similar extent in both accessions, albeit with a significant FDR in one accession, but not in the other. To avoid the selection of such genes, we used two more stringent approaches for the identification of genes that were clearly more (or only) affected by MD in one of the accessions. First, we selected genes that were DE in at least one accession and with a log_2_FC difference between two accessions under MD equal or larger than 1. As such, 138 genes were identified (Fig. 2D, Supplemental Table S5). Among these genes, 11 genes were DE in both accessions (but to a clearly different extent), and 79 and 48 genes were DE only in ICE163 or Yeg-1, respectively. Second, we identified twelve genes that showed a significant ecotype to treatment (E x T) interaction in the statistical model, with an FDR lower than 0.05 (Fig. 2E, Supplemental Fig. S3A). This set might include genes that were in the overlap of Figure 2A, but that were affected by MD to a much larger extent in one of the two accessions.

To further interpret the differences in transcriptional responses between ICE163 and Yeg-1, we combined the genes identified with these two methods, which resulted in a list of 143 genes (Supplemental Fig. S3B). A GO enrichment analysis was performed to gain further insight into the functional categories of these genes. The top-enriched categories included anthocyanin synthesis, JA signaling, response to ROS and response to abiotic stimulus (Supplemental Table S6). We selected genes based on their annotated functions in one of these GO categories (Fig. 2F, Supplemental Fig. S3C, Supplemental Table S7). It was interesting that more JA- and anthocyanin-related genes were induced in ICE163, while more ABA-related genes were induced in Yeg-1 by MD. With quantitative real-time PCR (qRT-PCR), we could confirm the observed differential expression of the majority of selected genes (Supplemental Fig. S4). As these results highlight that differences in drought sensitivity might be attributed to selective activation of anthocyanins, ROS, proline, and ABA mechanisms, we aimed at validating these transcriptomic findings by measuring these responses at physiological level. Wherever feasible, we included additional accessions that responded in a tolerant (EY15-2 and Sha) or sensitive (Oy-0 and ICE97) manner to the MD treatment.

### Anthocyanins Are Effective ROS Scavengers and Proline Acts as General Response Compound to Mild Drought

Drought is known to induce oxidative stress in plants by generating ROS (Miller et al., 2010). ROS can act as signaling molecules, but a strong accumulation can severely hamper plant growth and lead to cell death.

Our GO analysis of genes that were DE in ICE163 and Yeg-1 during MD showed a potential involvement of ROS, as illustrated by a differential regulation of *SODs* (Fig. 2F). Therefore, we examined the abundance of H_2_O_2_, one of the most prominent ROS, by performing a 3,3-diaminobenzidin (DAB) staining under WW and MD conditions. H_2_O_2_ levels, visualized as deep brown precipitates, were increased in the cotyledons of most accessions under MD, except in EY15-2 and ICE163 (Fig. 3A). However, no consistent dramatic differences between the tolerant and sensitive accessions could be observed.

**Figure 3.**
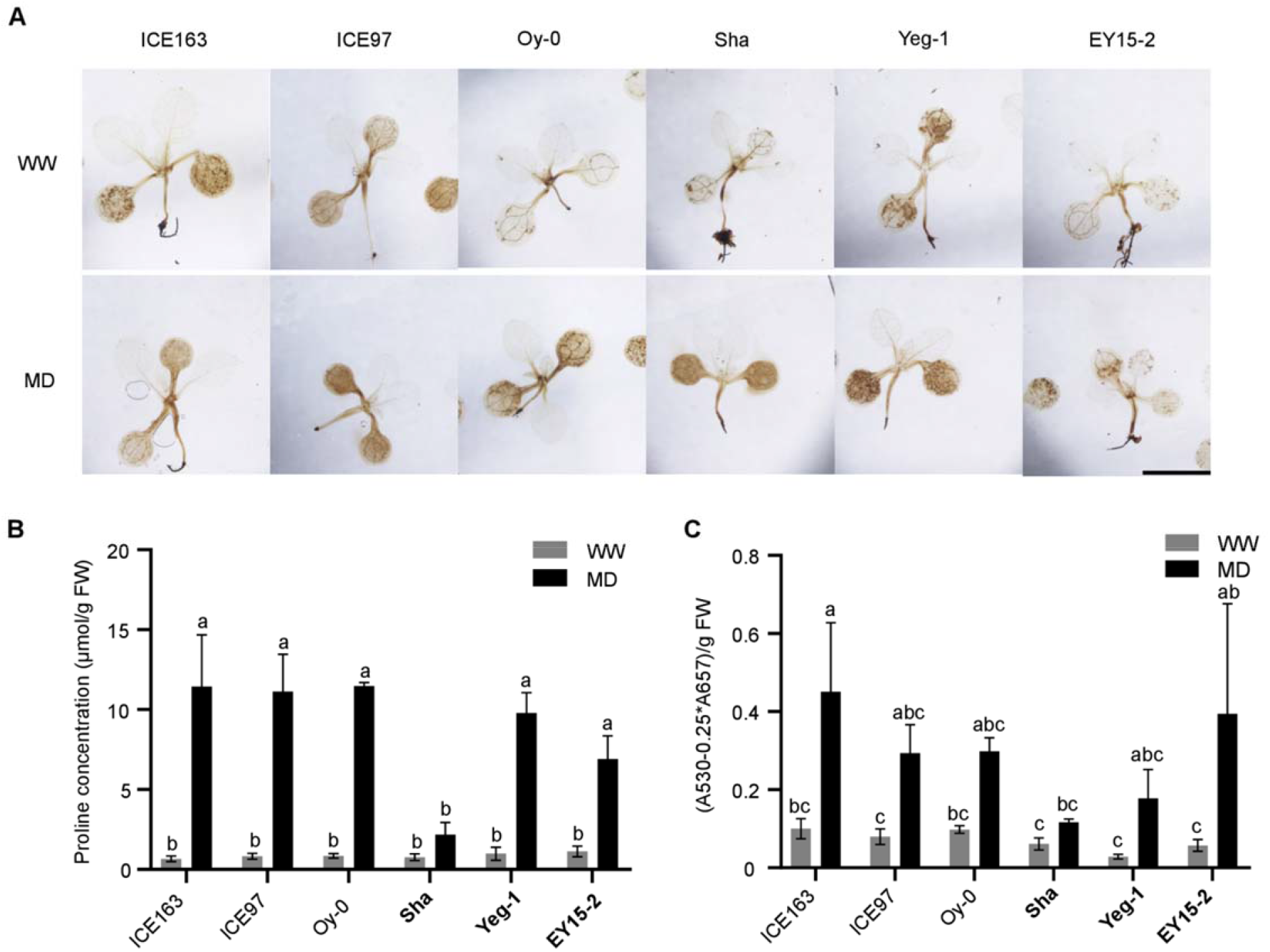
Proline, anthocyanins and ROS accumulation upon mild drought stress. **(A)** H_2_O_2_ accumulation in natural accessions seedlings detected by 3,3-diaminobenzidine (DAB) staining under well-watered (WW) and mild drought (MD) conditions. **(B)** Proline concentration of sensitive and tolerant accessions under WW and MD conditions at 11 DAS. **(C)** Anthocyanins concentration in sensitive and tolerant accessions under WW and MD conditions at 11 DAS. In B and C, error bars represent the SE, n = 3 biological repeats. Different letters above the error bar indicate statistical differences (p adj < 0.05; ANOVA, TukeyHSD). Tolerant accessions are in bold. FW, fresh weight.

To keep the homeostasis of ROS, plants have evolved complex enzymatic and non-enzymatic antioxidant systems, such as SODs and metabolites that act as antioxidants (Gill and Tuteja, 2010; Soares et al., 2019). Proline accumulation is known to play a positive role in abiotic stress (Bhaskara et al., 2015). In addition to proline, anthocyanin-related genes were overrepresented in our GO analysis. Because both proline and anthocyanins are able to scavenge ROS (Gould et al., 2002; Signorelli et al., 2014), we measured their abundance in seedlings at five days after water retention. Most accessions, except for Sha, accumulated proline to a similar level after the MD treatment (Fig. 3B). On the other hand, anthocyanins measurements revealed that the ecotypes that accumulated fewer H_2_O_2_, ICE163 and EY15-2, had a significant increase in anthocyanins content during MD (Fig. 3C). These results suggest that in our MD conditions, anthocyanins are effective in counteracting ROS, while proline acts as general drought response factor, both in sensitive and tolerant accessions.

### Tolerant Accessions Close Their Stomata More Efficiently

The closure of stomata under drought conditions is tightly controlled by ABA (Israelsson et al., 2006), one of the hormones that was highlighted by the differential transcriptome analysis in sensitive and tolerant accessions. We therefore measured the effect of drought on the stomatal closure of the tolerant and sensitive accessions at five days after water retention. In WW conditions, Oy-0 and ICE163 (drought-sensitive accessions) showed already a higher ratio of open stomata than ICE97 and the three tolerant accessions (Fig. 4, A and B). Under MD, stomatal opening was significantly reduced in all accessions (Fig. 4, A and B), but we found that the tolerant accessions had fewer open stomata than the sensitive accessions (Fig. 4B).

**Figure 4.**
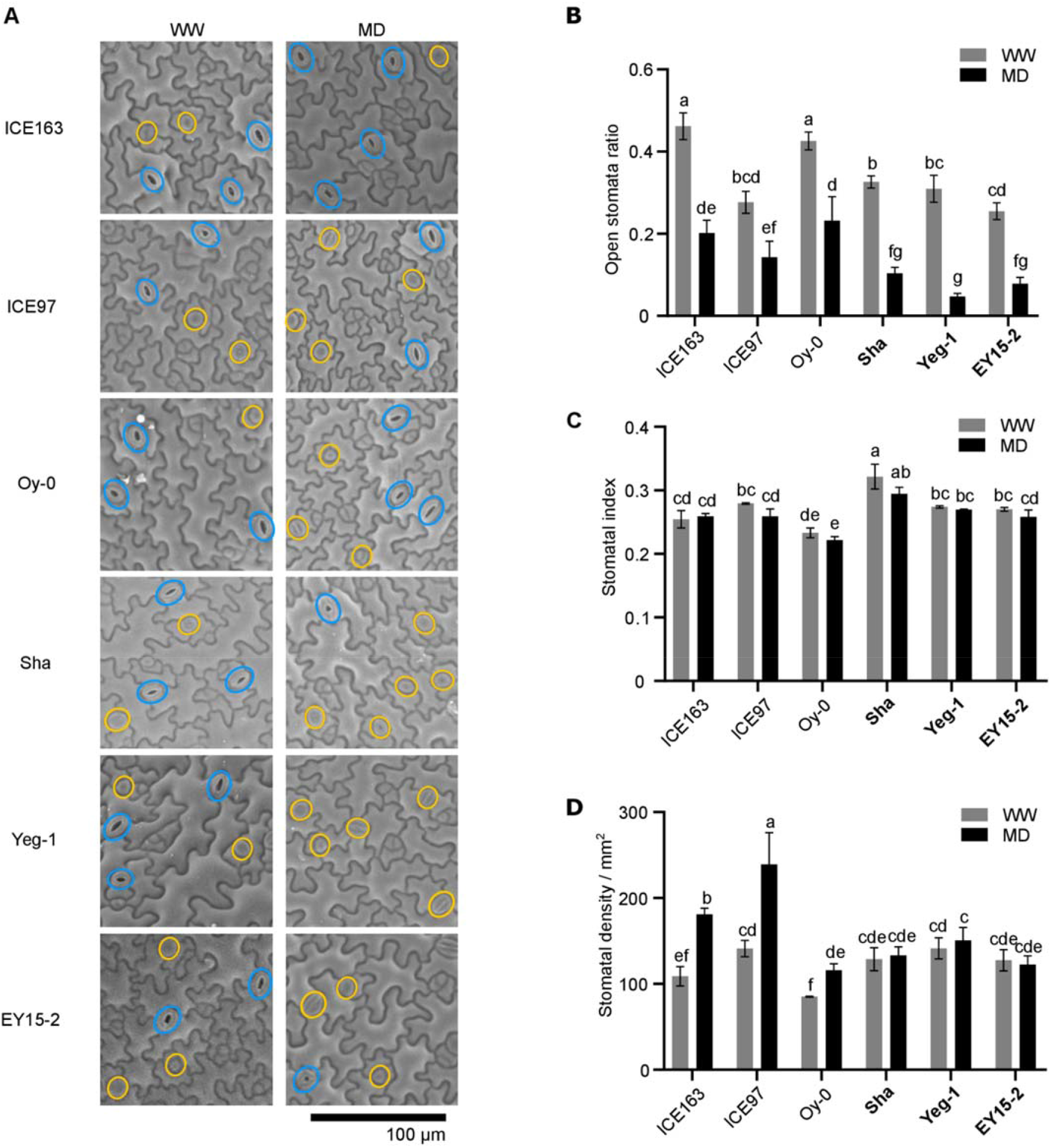
Stomatal opening, index and density after mild drought treatment. **(A)** Open and closed stomata in the first leaf pair under well-watered (WW) and mild drought (MD) conditions at 11 DAS. Blue circles indicate open stomata, and yellow circles indicate closed stomata. **(B)** Ratio of open stomata in sensitive and tolerant accessions under WW and MD conditions at 11 DAS. Error bars represent the SE, n = 3 biological repeats with ≥ 7 leaves per treatment, per ecotype, per repeat. **(C)** Stomatal index at 22 DAS. **(D)** Stomatal density at 22 DAS. Error bars represent the SE, n = 3 biological repeats, with 3 leaves per treatment, per ecotype, per repeat. Different letters above the error bars indicate statistical significances (p adj < 0.05; ANOVA, TukeyHSD). Tolerant accessions are in bold.

On top of the stomatal closure, it is reasonable to think that accessions with a higher number of stomata in a given leaf area can experience more stress when the soil water content decreases. Under MD conditions, plants with a lower stomatal density (SD, number of stomata per mm^2^) showed a lower transpiration and higher water use efficiency (Yoo et al., 2010; Franks et al., 2015). The SD of all sensitive and tolerant accessions was therefore analyzed at 22 DAS. Remarkably, the sensitive accessions ICE163 and ICE97 showed a significant increase in SD during MD treatment (Fig. 4D), whereas the SD was unaltered in the tolerant accessions.

The increase in SD in sensitive accessions could either result from an active decision to produce more stomata in the epidermis during drought, or from a maintained stomatal production while the growth of the leaf is arrested. To explore this, we calculated the stomatal index (SI, the number of stomata per total number of epidermal cells) at 22 DAS. From all accessions, Sha had the highest SI in both WW and MD conditions (32% and 29%, respectively), while Oy-0 had the lowest (23% and 22%, respectively) (Fig. 4C). However, we could not observe any significant effect on the SI by the MD treatment in all six accessions (Fig. 4C), suggesting that the development of stomata was not altered during drought.

Taken together, the tolerant accessions were found to close their stomata more efficiently than the sensitive accessions. The latter had a higher SD under MD stress, which is not resulting from an increased stomatal production but likely results from a decreased growth of non-stomatal cells.

### Drought-Tolerant Accessions Arrest Cell Proliferation but Maintain Cell Expansion

Our transcriptome data revealed that several cell wall-modifying enzymes, such as XYLOGLUCAN ENDOTRANSGLUCOSYLASE/HYDROLASES 19 (XTH19) and XTH24, were differently affected by drought in the sensitive vs. tolerant accession (Fig. 2F). Because leaf growth is strictly controlled by cell proliferation and cell expansion, we examined the effect of the MD treatment on these cellular characteristics in the selected sensitive and tolerant accessions. We found a significant reduction in average pavement cell numbers during drought in most accessions, except in EY15-2 (Fig. 5A), in which the final area of L3 was not significantly affected by drought (Fig. 1B). In all ecotypes, the cell number was reduced to a similar extent (Fig. 5A, Supplemental Fig. S5A). On the other hand, the average pavement cell area was dramatically reduced by the MD treatment in the sensitive accessions, whereas surprisingly no reduction could be observed in the tolerant accessions (Fig. 5B, Supplemental Fig. S5B). More specifically, the sensitive accessions showed an increased proportion of smaller cells or a decreased proportion of large pavement cells during MD treatment, but no significant differences could be observed in the tolerant accessions (Fig. 5C, Supplemental Fig. S6). These data indicate that the reduction in cell expansion is the major discriminating factor between tolerance and sensitivity to MD in these ecotypes.

**Figure 5.**
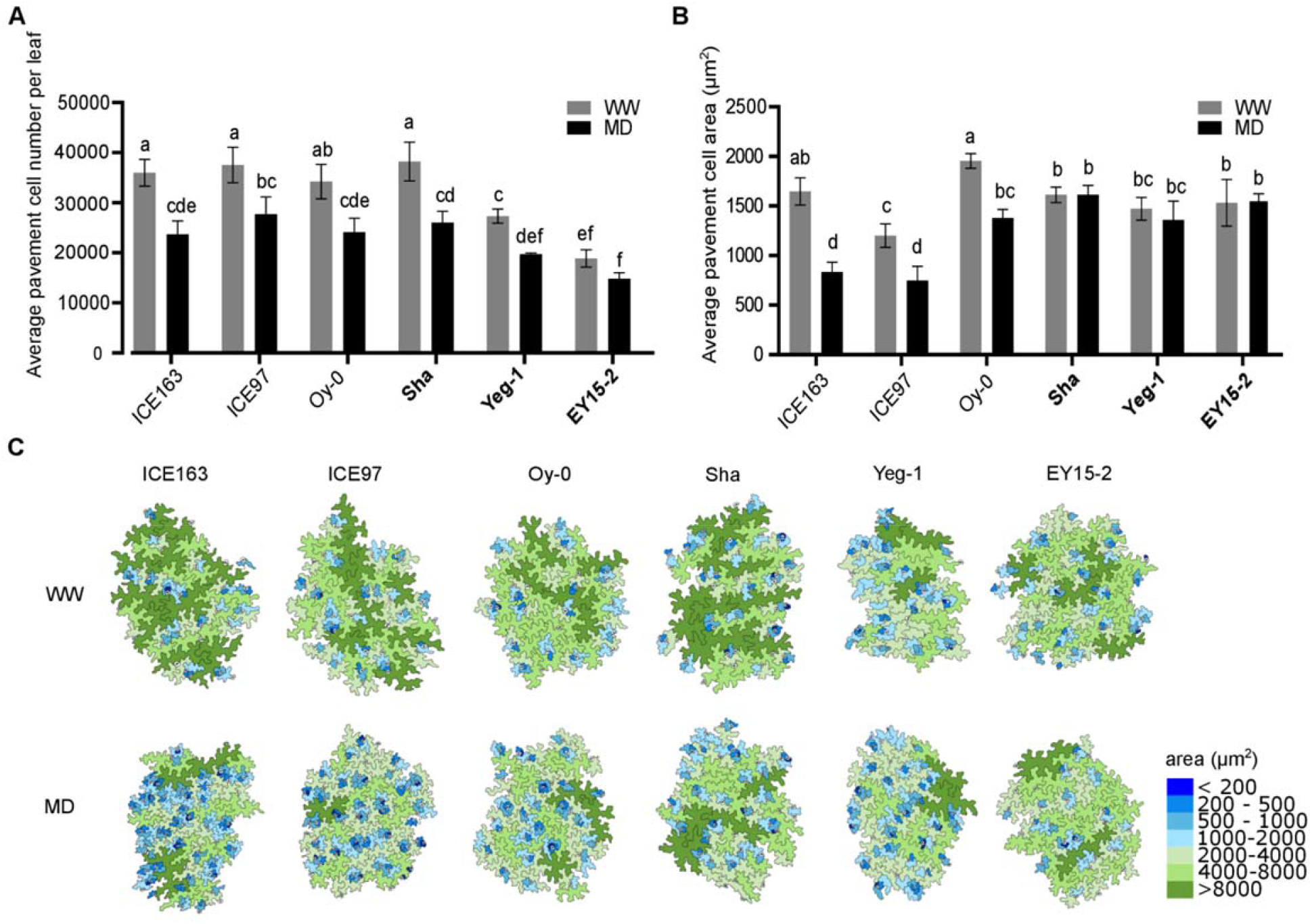
Mild drought differently affects pavement cell number and area in sensitive and tolerant accessions. **(A)** Average pavement cell number of leaf 3 (L3) under well-watered (WW) and mild drought (MD) conditions at 22 DAS. **(B)** Average pavement cell area of L3 under WW and MD conditions at 22 DAS. Error bars represent the SE, n = 3 biological repeats. Different letters above the error bar indicate statistical differences (p adj < 0.05; ANOVA, TukeyHSD). Tolerant accessions are in bold. **(C)** Representations of abaxial epidermal cell sizes of L3 of natural accessions at 22 DAS under WW and MD conditions.

## DISCUSSION

Arabidopsis natural accessions, which originated from different geographic regions, have evolved under different environmental conditions and therefore vary in many physiological traits, such as flowering time (Stinchcombe et al., 2004), circadian period length (Michael et al., 2003), hypocotyl length (Maloof et al., 2001) and root and leaf architecture (Pérez-Pérez et al., 2002; Chevalier et al., 2003; Rosas et al., 2013). Adapted to local habitats, they also show various tolerance levels to abiotic stresses, such as salt stress (Katori et al., 2010), drought stress (Des Marais et al., 2012), and low temperature (Hannah et al., 2006). In this study, we used fifteen natural accessions and indeed observed this diversity, with drought-triggered reductions in LA, ranging from 14% to 61%.

In Arabidopsis, molecular mechanisms contributing to stress tolerance can be found by performing transcriptome analysis of natural accessions, as the difference in tolerance between accessions can be achieved by the activation of specific stress-induced pathways. As such, mild drought induces a very limited number of common genes in mature leaves of different accessions (Des Marais et al., 2012). Similarly, our transcriptome analysis performed at early developmental stage identified much more accession-specific genes than those shared between the profiled tolerant (Yeg-1) and sensitive (ICE163) accession. In contrast, a previous report (Clauw et al., 2015) found 60 genes specifically DE between six tested accessions, whereas 354 genes were shared. However, the drought treatment had a similar effect on the reduction of L3 area in these six accessions, which explains the general molecular response (Clauw et al., 2015). Altogether, these three studies highlight an important specific response at the transcriptome level in accessions when they are exposed to a drought treatment that triggers accession-specific phenotypic effects.

From an agronomic point of view, it would be very interesting to reveal some of the discriminating factors that render plants either sensitive or tolerant to mild drought stress. Our study gives some insights into this by analyzing the differences between the sensitive accession, ICE163, and the tolerant accession, Yeg-1, at the transcriptomic, physiological, cellular and phenotypic level. First, some of the tested factors were equally affected by drought in the sensitive and the tolerant accessions. This was true for the accumulation of proline. High and equal proline accumulations were observed in most accessions, except in Sha, which is among the more tolerant accessions. Former research showed that proline accumulation is compromised in Sha because of defective alternative splicing of *DELTA1-PYRROLINE-5-CARBOXYLATE SYNTHASE 1* (*P5CS1*), which encodes a proline biosynthesis enzyme (Kesari et al., 2012). These observations confirmed previous findings that high accumulation levels of proline do not necessarily lead to an increased tolerance to drought stress (Kesari et al., 2012). At the phenotypic level, another trait that was affected by drought similarly in sensitive and tolerant accessions, is the reduction of cell division. Leaf growth is strictly coordinated by cell proliferation and cell expansion (Gonzalez et al., 2012), and reduction in cell division could be observed soon after exposure to mild drought conditions (Dubois et al., 2017). However, this reduction is not less affected in tolerant accessions. Together, these data suggest that sustaining cell division or proline accumulation during drought stress might not be a suitable strategy to maintain the growth of young leaves.

Second, our approach revealed accession-specific responses at transcriptome level that might, nevertheless, not result in important differences at physiological and tolerance level. For example, our transcriptome data suggested that the tolerant accession, Yeg-1, shuts down the JA and anthocyanin biosynthesis, while this was activated in the sensitive accession, ICE163. These results were surprising, as the levels of JA and anthocyanins were previously reported to be positively associated with the drought response (Sperdouli and Moustakas, 2012; Kazan, 2015). Indeed, application of exogenous JA can enhance drought tolerance in plants (Anjum et al., 2011) and overaccumulation of anthocyanins, as ROS scavenger, is positively associated with oxidative stress and drought tolerance (Nakabayashi et al., 2014). Several anthocyanin biosynthesis genes, such as *ANTHOCYANIDIN SYNTHASE* (*ANS*) and *DIHYDROFLAVONOL 4-REDUCTASE* (*DFR*), were upregulated while the biosynthesis repressor, *MYB-LIKE 2* (*MYBL2*), was downregulated in ICE163 by mild drought. Instead, JA signaling repressors, *JASMONATE-ZIM-DOMAIN PROTEIN 1* (*JAZ1*) and *JAZ7*, were upregulated in Yeg-1 by mild drought. In accordance with these expression data, we detected more anthocyanins accumulation in ICE163 compared to Yeg-1, under mild drought conditions. Antioxidant molecules, such as anthocyanins, can be efficient in keeping ROS homeostasis under mild drought. According to DAB staining, ICE163 accumulated less ROS in cotyledons than Yeg-1 upon mild drought treatment, which is opposite to anthocyanins accumulation levels. Both proline and anthocyanins have been reported to scavenge ROS under abiotic stress (Sperdouli and Moustakas, 2012; Signorelli et al., 2014) and we found that the lower levels of ROS in EY15-2 and ICE163 correlated with a higher anthocyanins content, suggesting an important role of anthocyanins in reducing ROS levels in these ecotypes. However, this ability to scavenge ROS more efficiently does not, or not sufficiently, contribute to the maintenance of growth under mild drought stress.

Then, which of the molecular and physiological strategies contributes to a significantly improved mild drought tolerance in young Arabidopsis leaves? One of the molecular pathways that was specifically induced in the tolerant Yeg-1 accession was ABA, one of the most important drought-responsive phytohormones regulating stomatal movements (Daszkowska-Golec and Szarejko, 2013). Yeg-1 showed a higher expression of the key biosynthesis gene *NINE-CIS-EPOXYCAROTENOID DIOXYGENASE 3* (*NCED3*) under mild drought, indicating a potentially higher ABA content, as overexpressing *NCED3* has been show to lead to an increase in ABA level, as well as an improvement in drought tolerance (Iuchi et al., 2001; Daszkowska-Golec and Szarejko, 2013). In accordance, we could also observe more closed stomata in Yeg-1 during mild drought. Stomatal closure in response to water deficits can limit photosynthesis (Chaves et al., 2003), but at the same time limit excessive decreases in water potential in plants (Van Houtte et al., 2013), thereby ensuring water demand for plant growth. Natural variations in stomatal response to closing stimuli have been observed (Aliniaeifard and van Meeteren, 2014). Here, we also observed various degrees of stomatal closure in natural accessions under mild drought, which is correlated with their mild drought tolerance ability.

On the other hand, ABA was previously also linked with cell expansion (Wu et al., 2018; Qiu et al., 2019) and, interestingly, this was another cellular parameter than we found to be differently affected in sensitive versus tolerant accessions. As mentioned earlier, cell expansion greatly contributes to leaf growth (Gonzalez et al., 2012) and is also decreased soon after exposure to mild drought conditions (Dubois et al., 2017). Genes from the *XTH* gene family encode cell wall-loosing enzymes, and some of them, such as *XTH19* and *XTH24*, are thought to positively influence cell elongation (Miedes et al., 2013; Lee et al., 2018). Under mild drought, *XTH19* was induced in Yeg-1, the tolerant accession, while *XTH24* was repressed in the sensitive accession ICE163, by mild drought. These differences in expression might explain the differences in cell area of L3 observed at the mature stage. Related to this, the maximal leaf expansion rate and the duration of leaf expansion were previously reported to be variable among natural accessions (Aguirrezabal et al., 2006). As such, An-1 is an accession that shows a prolonged leaf expansion upon drought, while Oy-0, a drought-sensitive accession, has a decreased duration of leaf 6 expansion (Aguirrezabal et al., 2006). We also observed that Oy-0 showed a decreased pavement cell area under mild drought, which might be caused by a shorter expansion duration, as reported.

In summary, our results indicate that different natural accessions in Arabidopsis use specific strategies to cope with mild drought, illustrated by the contrasting transcriptomic responses and the diverse physiological responses in tolerant and sensitive accessions. Interestingly, our analysis suggests that the ability to efficiently close stomata and maintain cell expansion under drought could be major determinants for the capacity to maintain leaf growth upon mild drought treatment. Our results further demonstrated that early responses in the transcriptome can reflect physiological responses at a later stage, such as *XTH* genes that can predict cell expansion phenotypes, or ABA biosynthesis genes that correlate with stomatal closure and improved tolerance at a later stage. Further research is needed to elucidate how tolerant accessions trigger more pronounced ABA responses, and to find key elements of this regulatory network. In addition, further examination of leaf growth, stomatal movement and cell expansion in mutants or overexpression lines of known ABA signaling pathway elements in different genetic backgrounds during mild drought would help to reveal common and accession-specific elements in mild drought tolerance.

## MATERIALS AND METHODS

### Plant Material, Growth Conditions and Mild Drought Treatment

Fifteen Arabidopsis natural accessions were selected to cover a broad range of genetic diversity of Arabidopsis. For the experiment of MD treatment in soil, seeds were sown in pots, which were filled up to 85 g ± 0.5 g of Saniflor soil (Van Isreal N.V., Geraardsbergen, Belgium) with an absolute water content of on average 70%. After 5 nights of stratification at 4°C in darkness, the pots were moved to a growth chamber (21°C and 16-h day/8-h night cycles). At 4 DAS, the pots were transferred to the automated phenotyping platform WIWAM (21°C and 16-h day/8-h night cycles) and randomized to homogenously mix the natural accessions and treatments. The controlled MD treatment was initiated at 6 DAS. Control-treatment pots were kept at 2.2 g water per gram dry soil. The MD-treated pots initially contained 2.2 g water per gram dry soil and their relative water content was further decreased to 1.2 g water per gram dry soil.

### Leaf Size Measurement

Leaf size measurements were performed on the third true leaf (L3) of the rosette at 22 DAS. The leaves were harvested, photographed, cleared in ethanol and subsequently mounted on microscopic slides in lactic acid. The leaf area was measured based on the pictures using ImageJ v1.45. Three biological repeats were performed for all accessions. Eight to twelve leaves were measured per treatment per repeat for all accessions, except for ICE197, for which four to nine leaves per treatment were measured.

### Statistical Analysis

All statistical analyses were performed with R, version 4.0.1 or Graphpad prism 8.4 (www.graphpad.com). ANOVA analyses were used for all leaf phenotypic and physiological data. Ecotype, treatment, repeat and their interactions were included as fixed effects of the model. Significant differences were assumed when with adjusted p-values lower than 0.05. The significance of the relative reduction in leaf area was determined by using the Phia package (https://cran.r-project.org/web/packages/phia/phia.pdf). The significant differences in relative frequencies of LN-transformed pavement cell area distribution was determined by ANOVA, using the Sidak’s multiple comparisons test.

### RNA-Sequencing Sampling

Eighty seedlings (distributed over nine to ten pots) were grown per ecotype per treatment per repeat. At 11 DAS, seedlings were harvested at 6 PM and stored in RNA-later (Ambion) solution. The third leaf was harvested by micro-dissecting, then pooled and flash frozen in liquid nitrogen. RNA extraction was done using Trizol (Invitrogen) combined with the RNeasy Mini Kit (Qiagen), following the manufacturer’s instructions. In total, three biological repeats were performed.

### RNA-Sequencing and Differential Expression Analysis

The sequencing was performed at the Nucleomics Core Facility (VIB, Leuven, Belgium, www.nucleomics.be), using a single-end mode with a read length of 75 bp on an Illumina NextSeq500. The quality control was done in the Galaxy platform with FastQC, alignment was done with Salmon. The Arabidopsis reference genome (TAIR10) was used.

The RNA-seq data was analyzed using R (version 3.6.0). Counts data of all splice variants were combined using the tximport package (https://bioconductor.org/packages/release/bioc/html/tximport.html). Then, the counts were normalized based on the library size of the samples. Genes that were detected less than three times with expression levels lower than 5 counts per million were removed. The new library was normalized by the TMM (trimmed mean of M values) method. A generalized linear model was applied with ecotype and treatment as factors using the glmQLFit function. Next, significant interactions were identified using the glmQLFTest function with the interaction term as a coefficient. The differential expression analysis of RNA-seq data was done with the EdgeR package (https://bioconductor.org/packages/release/bioc/html/edgeR.html) with R studio (Version 1.1.463). Differentially expressed genes of MD versus WW conditions for each ecotype were calculated using predefined contrasts. The cut-off of DE genes was set on a FDR < 0.05 and log_2_FC ≥ 0.6 or log_2_FC ≤ -0.6.

### Quantitative Real-Time PCR

The iScript cDNA Synthesis Kit (Bio-Rad) was used to synthesize the cDNA. The qRT-PCR was done on a LightCycler 480 (Roche Diagnostics) on 384-well plates with a LightCycler 480 SYBR Green I Master (Roche) according to the manufacturer’s instructions. Melting curves were analyzed to check the primer specificity. Normalization was done against the average of housekeeping genes AT1G13320, AT2G32170 and AT2G28390: ΔCt = Ct (gene) – Ct (mean [housekeeping genes]) and ΔΔCt = ΔCt (WW condition) – ΔCt (MD condition). Ct refers to the number of cycles at which the SYBR Green fluorescence reaches an arbitrary value during the exponential phase of amplification.

Primers were designed with the QuantPrime website (https://quantprime.mpimp-golm.mpg.de/main.php?page=home) and the conservation of their sequence in ICE163 and Yeg-1 was verified with POLYMORPH (http://polymorph.weigelworld.org/cgi-bin/webapp.cgi?page=show_aln;plugin=show_aln;project=MPICao2010). AT code, name of genes and primers used in this study are listed in Supplemental Table S8.

### Proline Content Measurements

Seedlings grown in control and drought conditions were harvested at 11 DAS. Free proline was measured by using the ninhydrin assay (Bates et al., 1973).

### 3,3-Diaminobenzidin (DAB) Staining

Seedlings were harvested at 11 DAS. For each treatment, eight seedlings were used for DAB staining per repeat. DAB was dissolved in a 10-mM PBS solution (1 mg/ml). Seedlings were submerged into the staining solution and vacuum-infiltrated three times for 5 min. Subsequently, the samples were incubated overnight in the dark. Chlorophyll was removed by adding ethanol to the samples and a boiling step for 10 min to obtain a better staining contrast. The cleared seedlings were transferred to 90% lactic acid and photographed.

### Anthocyanins Measurements

Seedlings were harvested, their fresh weight was measured and they were frozen in liquid nitrogen. The samples were homogenized and 1 ml of extraction buffer (45% methanol, 5% acetic acid) was added, followed by a thorough mixing. The mixtures were centrifuged at 12,000 x g for 5 min at room temperature. The supernatant was transferred to a new tube and centrifuged again for 5 min at room temperature. The absorbance of the supernatant was measured at 530 and 657 nm. The total anthocyanins content was calculated by (Abs 530 - 0.25 x Abs 657)/g fresh weight.

### Stomatal Closure

Imprints of the third leaf were made at 11 DAS at 6 PM. The nail polish copy of the imprint was analyzed using a tabletop scanning electron microscope (TM-1000, Hitachi). Seven to eight leaves were analyzed per accession per treatment in each biological repeat. Three random spots were checked for each leaf to calculate the average number of closed and open stomata.

### Cellular Analysis

For the cellular analysis, the total leaf blade area of cleared leaves was measured for at least eight representative leaves per repeat under a dark-field binocular microscope (MZ16, Leica). Abaxial epidermal cells in the center of the leaves were drawn using a microscope equipped with differential interference contrast optics (DM LB with 403 and 633 objectives; Leica) and a drawing tube. Scanned cell drawings were used to measure the pavement cell area as described before (Andriankaja et al., 2012). Cells were colored according to their area in Inkscape (https://inkscape.org/, RRID: <u>SCR_014479</u>).

## ACKNOWLEDGEMENTS

We would like to thank our colleagues of the Systems Biology of Yield group, and in particular Dr. Ting Li, for all constructive discussions and scientific advice. We also want to thank Véronique Storme for the statistical help on the article and Annick Bleys for her help in writing the manuscript.

